# Hippo-deficient cardiac fibroblasts differentiate into osteochondroprogenitors

**DOI:** 10.1101/2023.09.06.556593

**Authors:** Chang-Ru Tsai, Jong Kim, Xiao Li, Paulo Czarnewski, Rich Li, Fansen Meng, Mingjie Zheng, Xiaolei Zhao, Jeffrey Steimle, Francisco Grisanti, Jun Wang, Md. Abul Hassan Samee, James Martin

## Abstract

Cardiac fibrosis, a common pathophysiology associated with various heart diseases, occurs from the excess deposition of extracellular matrix (ECM)^1^. Cardiac fibroblasts (CFs) are the primary cells that produce, degrade, and remodel ECM during homeostasis and tissue repair^2^. Upon injury, CFs gain plasticity to differentiate into myofibroblasts^3^ and adipocyte-like^4,5^ and osteoblast-like^6^ cells, promoting fibrosis and impairing heart function^7^. How CFs maintain their cell state during homeostasis and adapt plasticity upon injury are not well defined. Recent studies have shown that Hippo signalling in CFs regulates cardiac fibrosis and inflammation^8–11^. Here, we used single-nucleus RNA sequencing (snRNA-seq) and spatially resolved transcriptomic profiling (ST) to investigate how the cell state was altered in the absence of Hippo signaling and how Hippo-deficient CFs interact with macrophages during cardiac fibrosis. We found that Hippo-deficient CFs differentiate into osteochondroprogenitors (OCPs), suggesting that Hippo restricts CF plasticity. Furthermore, Hippo-deficient CFs colocalized with macrophages, suggesting their intercellular communications. Indeed, we identified several ligand-receptor pairs between the Hippo-deficient CFs and macrophages. Blocking the Hippo-deficient CF-induced CSF1 signaling abolished macrophage expansion. Interestingly, blocking macrophage expansion also reduced OCP differentiation of Hippo-deficient CFs, indicating that macrophages promote CF plasticity.

## Introduction

Cardiac fibrosis, defined as the excess deposition of extracellular matrix (ECM), is associated with various type of heart diseases, such as myocardial infarction, hypertension, heart failure, and arrhythmia^1^. Cardiac fibroblasts (CFs) are the primary cell type that produces, degrades, and remodels the ECM^3^. Previously, different cell states of CFs were identified during homeostasis and tissue repair through genetic lineage tracing and single-cell transcriptomic profiling studies^7^. Nonetheless, the molecular and cellular mechanisms of these cell state transitions remain poorly defined. Moreover, how non-CFs interact with CFs and affect CF cell state alterations during cardiac fibrosis remain to be explored.

The Hippo pathway is a highly conserved signaling pathway that regulates cell differentiation, cell polarity and the balance between cell proliferation and cell death^12,13^. The core Hippo kinase cascade composed of MST1/2 and LATS1/2 phosphorylate and inhibit the downstream transcription co-activator, YAP and TAZ. In turn, YAP and TAZ bind to 14-3-3 and are retained in the cytoplasm. In the absence of Hippo signaling, YAP and TAZ enter the nucleus and bind to transcription factors, such as TEA domain family members (TEAD1-4), to promote downstream gene expression. Inhibition of Hippo signaling in cardiomyocytes promote regeneration in mice^14^ and porcine^15^. In contrast, inhibition of Hippo pathway in CFs promote cardiac fibrosis and inflammation^8^. Furthermore, loss of YAP and TAZ reduced fibrosis and inflammation upon cardiac injuries^9–11^.

In this study, we found that deleting the Hippo core kinase Lats1/2 in CFs promoted the differentiation of CFs into osteochondroprogenitors (OCP). In addition, we found that Hippo-deficient CFs secreted colony stimulating factor 1 (CSF1) to promote macrophage expansion. Given that blocking macrophage expansion reduced fibrosis, macrophage expansion in the context of Hippo-deficient CFs is profibrotic. Interestingly, OCP differentiation is also partially dependent on macrophage, indicating that macrophages have a critical role in the CF cell state transition during fibrosis.

## Results

### Hippo-deficient cardiac fibroblasts induce arrhythmia, inflammation, and fibrosis

In human mutations in genes encoding Hippo pathway components are associated with heart defects and diseases (Fig. 1A). Single-nucleotide polymorphisms (SNPs) in *MOB1B* and near *TEAD4* and *YWHAE* (14-3-3ε) have been linked to atrial fibrillation (Table 1), whereas genome-wide association studies (GWAS) showed that variants in Hippo pathway components genes *MST1/2, LATS1/2, YAP*, and *TAZ* are associated with various septal and valve defects (Table 1). To investigate the detailed molecular and cellular mechanisms of Hippo signaling in CFs, we used an inducible Cre transgenic mouse line (*Tcf21^iCre,^*^16^) to generate mice with the knockout of *Lats1/2* specifically in mouse CFs, hereafter referred to as *Lats1/2^ΔCF^* mice. At the indicated timepoints, we performed telemetry, histology, single nucleus RNA sequencing (snRNA-seq) and spatial transcriptomics (ST) analyses (Fig. 1B). Electrocardiographic analysis of *Lats1/2^ΔCF^* mice showed an irregular RR interval (Fig. 1C,D), an inverted P-wave (Extended Data Fig. 1A), an increased PP interval (Extended Data Fig. 1B) and a slower heart rate (Extended Data Fig. 1C). These finding in mice could be linked to the arrhythmia phenotype observed in patients with genetic variants of Hippo pathway components (Fig. 1A). Two weeks after Cre activation, histology analysis of control and *Lats1/2^ΔCF^* mice revealed significant hyperplasia of the right atrium (RA) in *Lats1/2^ΔCF^* mice (Extended Data Fig. 1D). To better examine fibrosis, we stained collagen with Sirius red staining and observed substantially increased collagen signals in the RA (Fig. 1E,F). We hypothesized that Yap and Taz were the critical downstream targets of Lats1/2 in this context and that deleting *Yap* and *Taz* would suppress the phenotypes induced by Hippo-deficient CFs. Indeed, the knockout Yap/Taz in *Lats1/2^ΔCF^* mice substantially reduced fibrosis (Fig. 1E,F) and hyperplasia (Extended Data Fig. 1D, lower panels) in the RA. Interestingly, neither hyperplasia (Extended Data Fig. 1D, right panel) nor fibrosis (Extended Data Fig. 1E) was observed in the left atria (LA) of *Lats1/2^ΔCF^* mice.

**Fig. 1.**
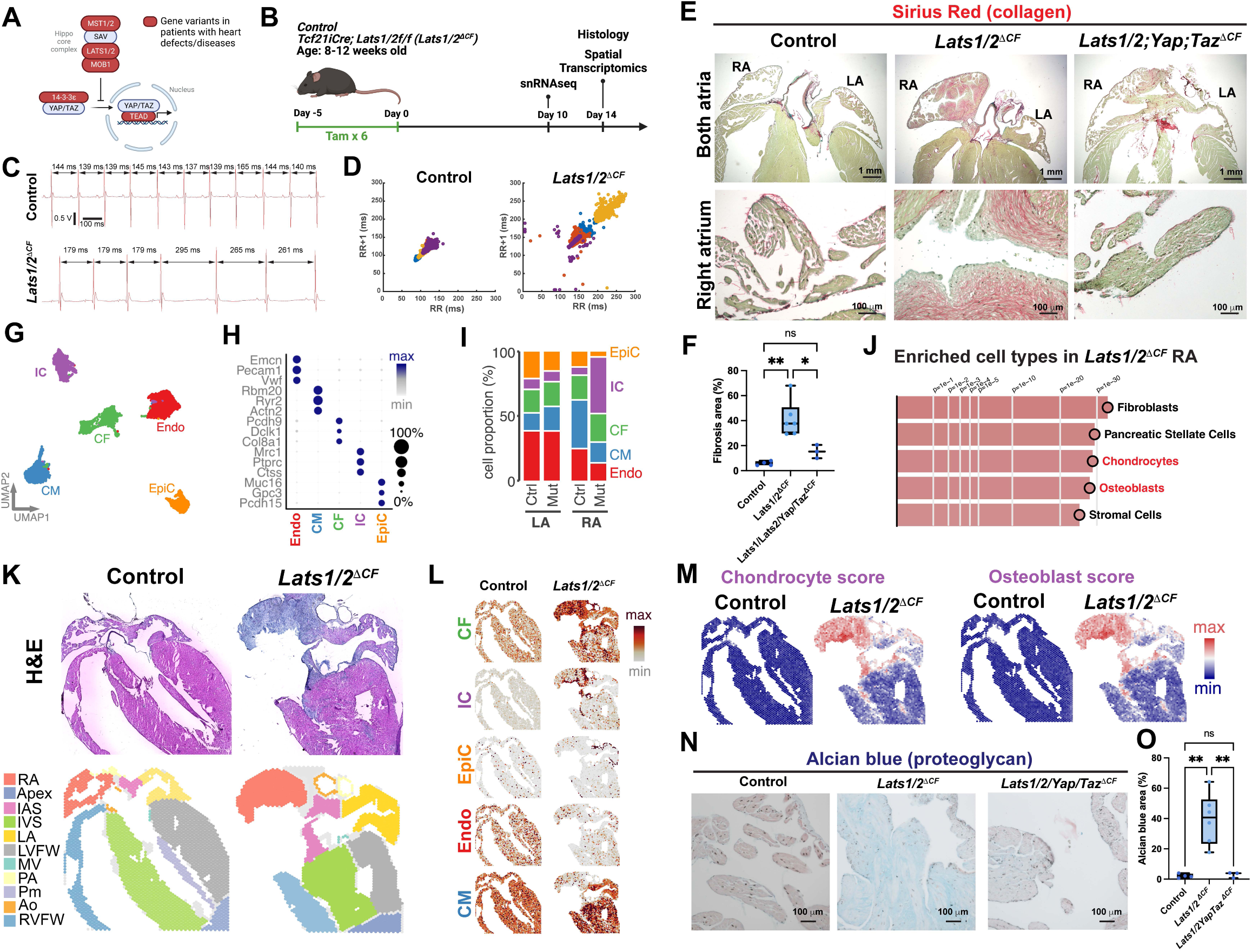
Hippo-deficient cardiac fibroblasts promote arrhythmia, inflammation, and fibrosis. (A) Diagram of the core Hippo pathway components. Genes associated with heart defects and diseases are highlighted in red. (B) Experimental scheme. (C-D) Electrocardiograph from control and *Lats1/2^ΔCF^*mice. (C) A representative example of arrhythmia in a *Lats1/2^ΔCF^* mouse three weeks after Cre activation. Double-headed arrows indicate RR intervals. (D) Representative Poincaré plots showing beat-to-beat RR interval variability from control (n=4) and *Lats1/2^ΔCF^* (n=4) mice. Each color represents individual RR/RR+1 intervals from a different mouse. (E) Sirius red staining (collagen) of heart sections from control, *Lats1/2^ΔCF^*, and *Lats1/2YapTaz^ΔCF^* mice collected two weeks after Cre activation. Upper panels show zoomed-out images. Lower panels show zoomed-in images. Right atrium, RA; left atrium, LA. (F) The percent fibrotic area defined as the collagen area over total tissue area for control (n=4), *Lats1/2^ΔCF^*(n=6), and *Lats1/2YapTaz^ΔCF^*(n=3) mice. One-way ANOVA, multiple comparisons. (G-I) Single nucleus RNA sequencing (snRNA-seq) analysis showing the (G) uniform manifold approximation and projection (UMAP) of major cell types in control and *Lats1/2^ΔCF^* atria. Major cell types included epicardial cells (EpiC), cardiac fibroblasts (CF), endothelial cells (Endo), cardiomyocytes (CM), and immune cell (IC). (H) Differentially expressed genes in different cell types. (I) Cell type composition in each sample. (J-M) Spatial transcriptomics (ST) data. (K) Cryosections of control and *Lats1/2^ΔCF^* hearts collected two weeks after Cre activation. Heart sections are shown stained with hematoxylin and eosin and anatomically annotated. IAS, inter-atrial septum; IVS, inter-ventricle septum. LVFW, left ventricular free wall; RVFW, right ventricular free wall; MV, mitral valve. PA, pulmonary artery; Pm, papillary muscle; Ao, aorta. (L) Deconvolutions of major cell types in the ST data. (K) ST data showing enriched cell types in the RA of *Lats1/2^ΔCF^* mice compared with control mice. (M) ST data-derived chondrocyte and osteoblast scores in control and *Lats1/2^ΔCF^* hearts. (N) Alcian blue (proteoglycan) staining of RA sections from control (n=4), *Lats1/2^ΔCF^* (n=6), and *Lats1/2YapTaz^ΔCF^* (n=3) mouse hearts. (O) Percentages of Alcian blue area was defined as Alcian blue area over the total tissue area. Welch’s test. (F,O) ns, not significant. *, P<0.05. **, P<0.01.

**Table. 1.** Genetic variants of Hippo pathway components associated with heart defects and diseases.

Next, to further rule out the possibility that the phenotype in the RA is not due to higher *Tcf21iCre* activity or more fibroblasts in the RA, we utilized Pdgfra-H2B-EGFP knock-in mice (*Pdgfra^EGFP^*) which all *Pdgfra*-positive fibroblasts are marked by nuclear GFP. We crossed *Pdgfra^EGFP^* mice with *Tcf21^iCre^; Rosa26^Ai-9^* (Cre lineage was labeled by red fluorescent protein, Tomato). The pups from the resulting litter with all three transgenic alleles were treated with tamoxifen to activate Cre so that the Cre positive cells would be Tomato-positive, and all of the CFs would be GFP-positive. We found that the CF (GFP-positive) composition was comparable between the LA and RA (∼25%, Extended Data Fig. 2A,B). *Tcf21^iCre^* efficiency in CFs, which was measured as the number of Tomato and GFP double-positive cells over the number of total GFP-positive cells were also similar between the LA and RA (Extended Data Fig. 2A,C).

To further determine whether the *Lats1* and *Lats2* exons flanked by the LoxP sites were indeed recombined and deleted in both atria, we performed quantitative real-time PCR (qPCR) to detect the recombination events. We designed primers flanking the deleted exon (recombination primer sets) as well as a nearby non-deleted exon as internal control (internal control primer sets) (Extended Data Fig. 2D) and used the cycle threshold (Ct) number from the qPCR reactions to assess the DNA input and the recombination events in the LA and RA of *Lats1/2^ΔCF^* mice. For the internal *Lats1* and *Lats2* primer set, no significant difference was observed between the LA and RA of *Lats1/2^ΔCF^*mice (Extended Data Fig. 2E,F), indicating that the amount of input DNA template was comparable. For the recombination *Lats1* and *Lats2* primer sets, no significant difference was observed between LA and RA of *Lats1/2^ΔCF^* mice (Extended Data Fig. 2G,H), indicating that the recombination rate in *Lats1* and *Lats2* flox loci was comparable between the LA and RA. Therefore, we concluded that the weaker phenotype in the *Lats1/2^ΔCF^* LA was not due to low Cre activity. The cause(s) of phenotypic difference between *Lats1/2^ΔCF^* LA and RA is currently under investigation for another study. Hereafter we will focus on the RA.

To understand the detailed mechanisms underlying the hyperplasia and fibrosis induced by Hippo-deficient CFs at the single cell level, we isolated nuclei from the LA and RA of control and *Lats1/2^ΔCF^* mice 10 days after Cre activation and performed snRNAseq (Fig. 1B). In an unbiased cell clustering analysis, five major clusters were identified (Fig. 1G). Based on differential gene expression (Fig. 1H) and cell type-specific markers, we identified the clusters as cardiomyocytes (CMs), CFs, epicardial cells (EpiCs), endothelial cells (ECs), and immune cells (ICs). By comparing cell type composition between samples, we found that ICs were substantially increased in the RA of *Lats1/2^ΔCF^* mice compared with control mice (Fig. 1I), which was consistent with our previous finding that Hippo-deficient CFs promote inflammation in the ventricle^8^. Similar to the histological analysis (Extended Data Fig. 1D,E), no significant alteration in cell type composition was observed between the LA of control and *Lats1/2^ΔCF^* mice (Fig. 1I).

To investigate how Hippo-deficient CFs interact with surrounding cells and affect tissue architecture, we performed ST analysis of control and *Lats1/2^ΔCF^* hearts two weeks after Cre activation (Fig. 1K). Hematoxylin and eosin staining (Fig. 1K, upper panel) of the ST data was used to annotate the anatomy of the heart (Fig. 1K, lower panel). To integrate the snRNA-seq and ST datasets, we deconvoluted the ST data by using major cell type markers identified in the snRNAseq dataset. We found that CFs and ICs were significantly increased in the RA of *Lats1/2^ΔCF^* mice compared with control mice, whereas CMs were reduced (Fig. 1L). To compare the changes in cell type composition in the RA between control and *Lats1/2^ΔCF^* mice, we performed cell type enrichment analysis and found fibroblasts were enriched in the RA of *Lats1/2^ΔCF^* mice compared to control mice (Fig. 1J). Interestingly, we also found that chondrocytes and osteoblasts were enriched in the RA of *Lats1/2^ΔCF^*. Indeed, in the ST data, the highest expression of chondrocyte and osteoblast markers was observed in the RA of *Lats1/2^ΔCF^* mice (Table 2, Fig. 1M). To understand which cell types gained chondrocyte and osteoblast features, we scored osteoblast and chondrocyte markers in the snRNAseq dataset and found that CFs had the highest scores (Extended Data Fig. 3A,B), suggesting that CFs may differentiate into chondrocyte-like and osteoblast-like cells. Because most of the enrich genes for osteoblasts and chondrocytes were overlapping (Fig. S3C), the possibility remains that Hippo-deficient CFs differentiate into the osteochondroprogenitor (OCP) cell state. To further examine the presence of OCP cells, we performed Alcian blue staining, which is a typically used to detect chondrocyte-secreted proteoglycans in cartilage. Increased Alcian blue was observed in the RA of *Lats1/2^ΔCF^* mice compared with control mice (Fig. 1N,O). Moreover, increased Alcian blue staining signal was reduced when *Lats1/Lats2/Yap/Taz* were simultaneously deleted (Fig. 1N,O), indicating a dependence on YAP/TAZ and that YAP and TAZ are the critical downstream targets in this context. Furthermore, these results suggest that Hippo-deficient cardiac fibroblasts differentiate into OCP cells.

**Table. 2.** ST data-derived lists of chondrocyte and osteoblast markers enriched in the right atria of *Lats1/2^ΔCF^* mice.

### Hippo-deficient cardiac fibroblasts differentiate into osteochondroprogenitors

To better understand how the Hippo pathway regulates CFs in the RA, we pooled and reclustered CFs from the RA of control and *Lats1/2^ΔCF^*mice. We identified three major subclusters named CF1-3 (Fig. 2A). By comparing the differential gene expression of these clusters and sample source (Fig. 2B), we found that CF1 primarily consisted of CFs from the RA of control mice. Classical CF markers, such as Gelsolin (Gsn), were highly expressed (Fig. 2C, left upper panel), representing the ground state of CFs during homeostasis. CF3 mostly consisted of CFs from the RA of *Lats1/2^ΔCF^* mice. Canonical YAP targets, such as *Amotl2* (Fig. 2C, left lower panel) and *Vgll3* (Extended Data Fig. 4A, left panel) were highly expressed in CF3, representing *Lats1/2* conditional knockout (*Lats1/*2CKO) CFs. CF3 also had the highest YAP score (Fig. 2D), determined by measuring the expression of 71 previously identified YAP targets^8,17^ (Table 3). Along the same lines, ST data showed the higher expression of Amotl2 (Fig. 2C, right lower panel) and Vgll3 (Extended Data Fig. 4A, right panel), and a higher Yap score (Fig. 2G, upper panel) in the RA of *Lats1/2^ΔCF^* mice than in that of control mice. To validate that CF3 was indeed composed of *Lats1/*2CKO CFs, we performed in situ hybridization to detect *Amotl2* and *Vgll3* transcripts and costained the cells for GFP to mark *Tcf21^iCre^* lineage cells in the RA of control and *Lats1/2^ΔCF^* mice. We found that *Amotl2* and *Vgll3* were specifically upregulated in the RA of *Lats1/*2*^ΔCF^*mice but not of control mice and were mostly GFP-positive (Extended Data Fig. 4B, red), supporting that *Tcf21^iCre^* lineage cells marked by GFP represented the CF3 subcluster. Because *Tcf21^iCre^* efficiency was only approximately 40% in the RA (Extended Data Fig. 2C), we reasoned that CF2 cells from the RA of *Lats1/*2*^ΔCF^* mice (Fig. 2A,B) with a lower YAP score than CF3 cells were likely to have escaped from *Tcf21^iCre^* recombination and *Lats1/2* deletion (Fig. 2D). Although CF2 cells were likely to be wild-type cells in the *Lats1/*2*^ΔCF^* RA, they expressed higher level of *Col1a1, Acta2,* and *Tagln2* (Extended Data Fig. 4C) than CF1 cells did. This suggested that CF2 cells were activated, which may affect fibrosis in this context.

**Fig. 2.**
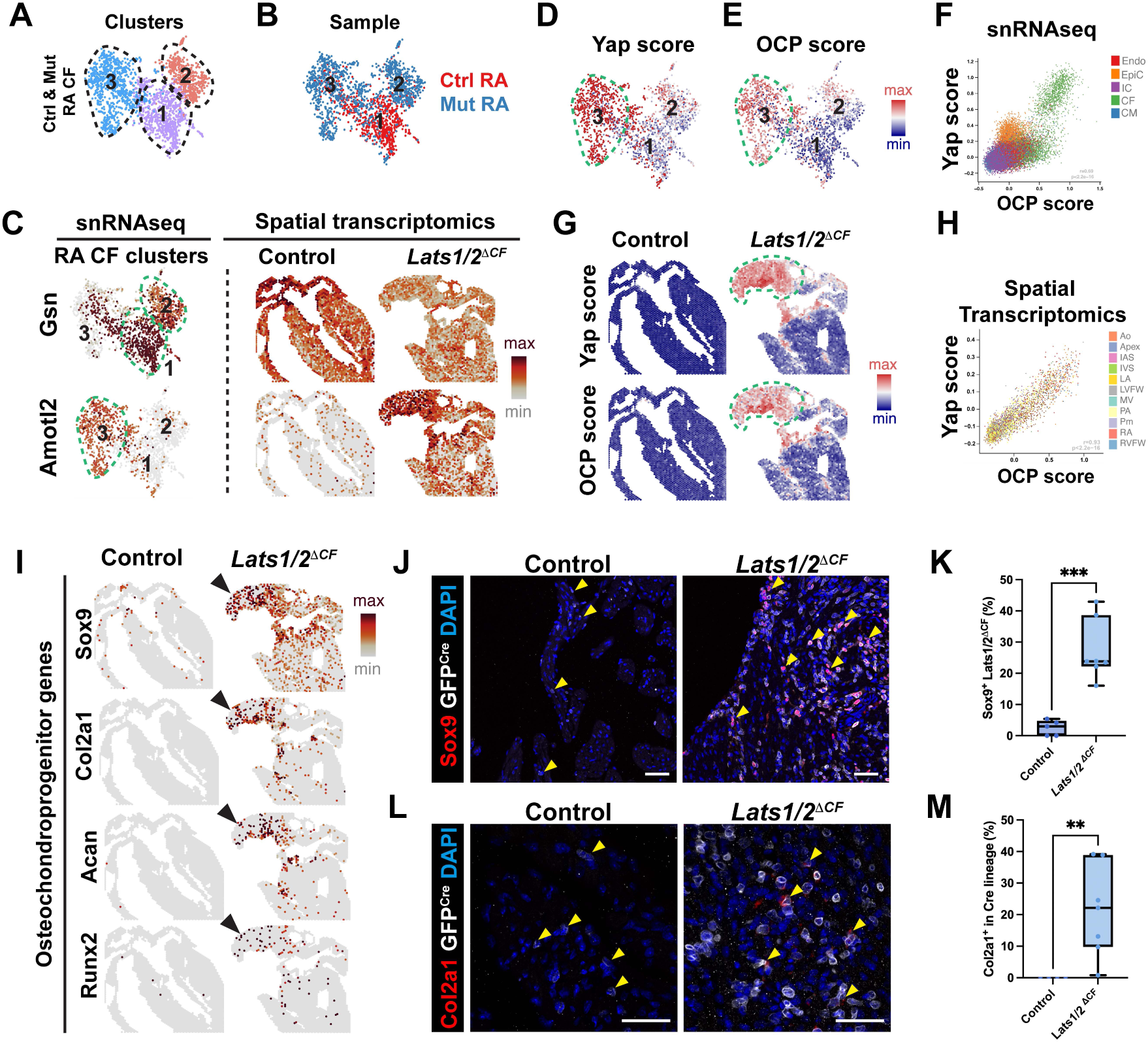
Differentiation of right atrial Hippo-deficient cardiac fibroblasts into osteochondroprogenitors. (A) UMAP of CF subclusters from the RA of control and *Lats1/2^ΔCF^*mice. (B) Sample source of CF subclusters in the RA. (C) Expression of Gsn (upper panels), a resting CF marker, and Amotl2 (bottom panels), a classical YAP target, in snRNAseq data (left panels) and ST data (right panels). Expression of YAP targets (YAP score) (D) and OCP markers (OCP score) (E) in the snRNA-seq data and their correlation (F). Expression of Yap targets (YAP score) and OCP markers (OCP score) (G) in ST data from control and *Lats1/2^ΔCF^* hearts and their correlation (H). (I) OCP marker expression in control and *Lats1/2^ΔCF^* hearts. (J) RA sections of control and *Lats1/2^ΔCF^* mouse hearts stained with antibodies against SOX9 (red), which is an OCP marker, and GFP (white), which is used to label Cre lineage cells. DAPI (blue) was used to stain nuclei. Scale bar, 50 μm. (K) Percentage of SOX9-positive cells in *Tcf21^iCre^* lineage cells (green). Control, n=5. *Lats1/2^ΔCF^*, n=7. Welch’s test was used for the comparison. (L) In situ hybridization of *Col2a1* mRNA transcript (an OCP marker, red) and immunostaining of GFP (Cre lineage, white) in the RA of control and *Lats1/2^ΔCF^* mice. DAPI (blue) was used to stain nuclei. Scale bar, 50 μm. (M) Percentage of *Col2a1*-positive cells in the *Tcf21^iCre^* lineage cells in control (n=4) and *Lats1/2^ΔCF^* mouse hearts (n=5). Mann-Whitney test was used for comparisons. (K,M) **, P<0.01; ***, P<0.001.

**Table. 3.** Lists of YAP targets used to measure the YAP score.

Next, to examine which CF subclusters differentiated into OCPs, we scored the expression of chondrocyte and osteoblast markers amount the three CF subclusters. CF3 cells had the highest OCP score (Fig. 2E) and YAP score (Fig. 2D). Indeed, in the snRNA-seq (Fig. 2F) and ST (Fig. 2G) datasets, the OCP score and YAP score were highly correlated, particularly in the RA of *Lats1/*2*^ΔCF^* mice. Furthermore, the expression of OCP markers, including *Sox9, Col2a1, Col11a1, Col12a1, Acan,* and *Runx2*, was increased in the RA of *Lats1/*2*^ΔCF^* mice than the control (Fig. 2I and Extended Data Fig. 4D). To validate the snRNAseq and ST data, we performed immunostaining with antibodies against Sox9 and GFP and found that SOX9 protein expression was increased in *Tcf21^iCre^* lineage cells (labeled by GFP) in the RA of *Lats1/*2*^ΔCF^* mice compared with that of control mice (Fig. 2J,K). Moreover, >90% SOX9-positive cells were also GFP-positive (Extended Data Fig. 4E), indicating that Yap activation autonomously promotes the upregulation of SOX9 protein expression. In situ hybridization data showed that the expression of *Col2a1*, a critical downstream target of SOX9^18^, was also increased in *Lats1/2*CKO CFs (Fig. 2L,M). These results suggested that *Lats1/2* are critical for suppressing OCP differentiation in CFs during homeostasis.

### Hippo-deficient cardiac fibroblasts secrete CSF1 to promote macrophage expansion

To determine which types of cells interact with Hippo-deficient CFs, we first deconvoluted the ST data from *Lats1/2*CKO CFs using CF3 markers identified in the snRNA-seq analysis. The resulting cells, named YAP^High^ CFs, were highly enriched in the RA of *Lats1/*2*^ΔCF^* mice but not control mice (Fig. 3A, right panel). In contrast, YAP^Low^ CFs, as deconvoluted using CF1/2 markers, were present in the RA of both control and *Lats1/*2*^ΔCF^* mice (Fig. 3A, left panel). Using the deconvoluted ST data, we compared major cell types (Fig. 1L, middle panel) and found that ICs and YAP^High^ CFs (Fig. 3A, right panel) showed similar pattern and were indeed colocalized (Fig. 3B). Most cells in the IC cluster were macrophages as they expressed high level of macrophage markers, Cx3cr1, Csf1r, Itgb9 (data not shown).

**Fig. 3.**
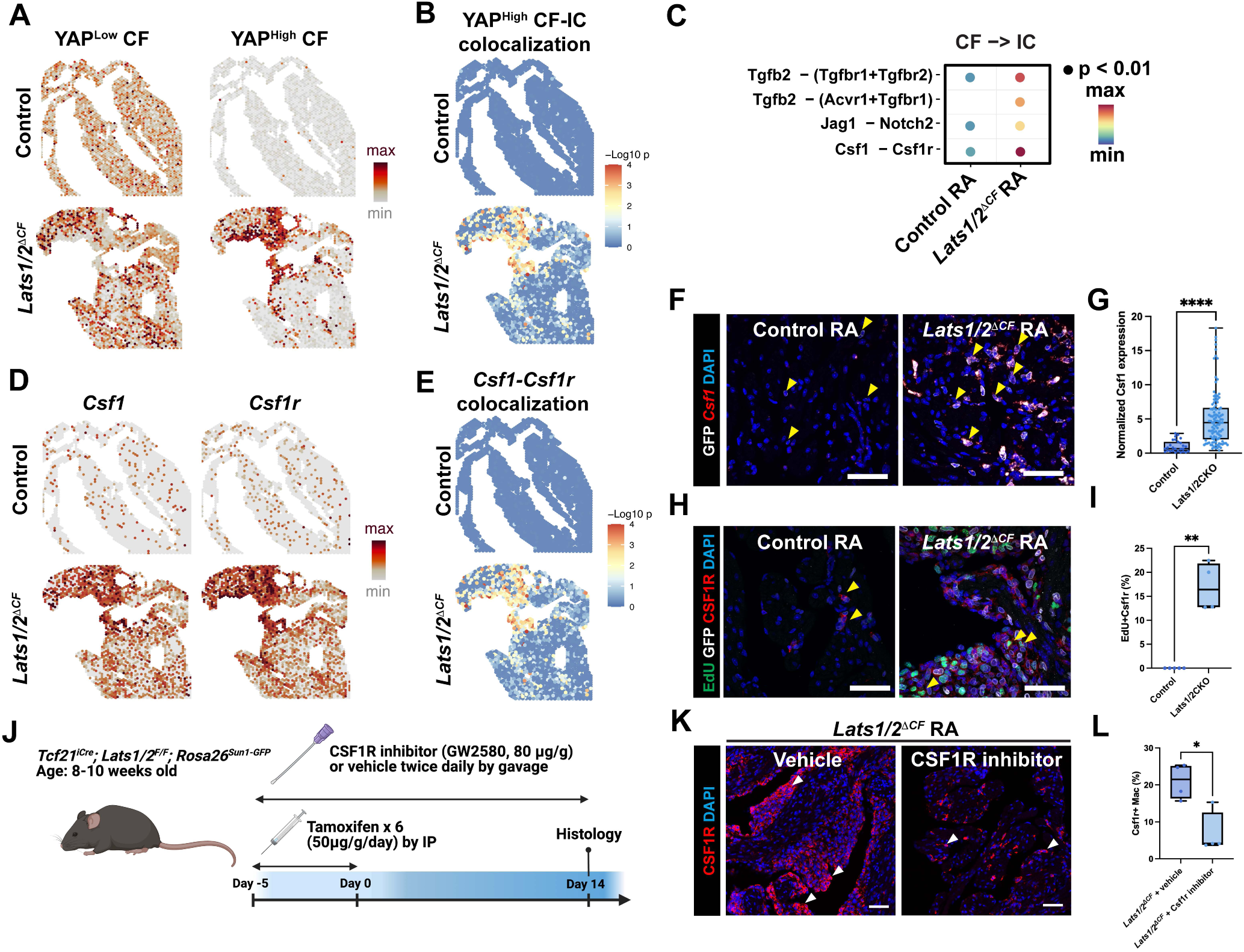
Hippo-deficient cardiac fibroblasts secrete CSF1 to promote macrophage expansion. (A) CFs with a low YAP score (YAP^low^ CF), and CFs with a high YAP score (YAP^high^ CF), were deconvoluted in the ST data by using their cell markers. (B) YAP^high^ CF and IC colocalization in the ST data. (C) Ligand-receptor pairs between CFs and ICs in the RA of control and *Lats1/2^ΔCF^* mice. (C) ST data showing *Csf1* and *Csf1r* expression and colocalization (E) in control and *Lats1/2^ΔCF^* mouse hearts. (F) RNAscope detecting *Csf1* transcripts (red) in *Tcf21^iCre^* lineage cells labeled by a GFP reporter (white). DAPI (blue), was used to stain nuclei. Yellow arrowheads point to examples of GFP^+^ cells. Scale bar, 50 μm. (G) *Csf1* expression intensity in *Tcf21^iCre^*lineage cells in the RA of control and *Lats1/2^ΔCF^* mouse heart. Each dot represents each *Tcf21^iCre^* lineage cell (27 cells from four different control mice and 107 cells from four different *Lats1/2^ΔCF^* mice). The Mann-Whitney test was used for comparisons. (H) Immunostaining of *Tcf21^iCre^* lineage cells and macrophages using GFP (white) and CSF1R antibodies (red), respectively, in the control and *Lats1/2^ΔCF^* RA with EdU labeling to mark proliferating cells (green). DAPI (blue) was used to stain nuclei. Yellow arrowheads point to examples of CSF1R-positive macrophages. Scale bar, 50 μm. (I) Percentage of EdU-positive *Tcf21^iCre^* lineage cells in the RA of control (n=5) and *Lats1/2^ΔCF^*hearts. The Mann-Whitney test was used for comparisons. (J) Experimental scheme. (K) CSF1R antibody (red) was used to label macrophages in the *Lats1/2^ΔCF^* RA treated with Csf1r inhibitor or vehicle. DAPI (blue) was used to stain nuclei. White arrowheads point to examples of CSF1R-positive macrophages. Scale bar, 50 μm. (L) Percentages of CSF1R-positive macrophages among all cells (DAPI) in the RA of *Lats1/2^ΔCF^* mice treated with CSF1R inhibitor or vehicle. n=4 per group. The Mann-Whitney test was used for comparisons. (G,I,L) *, P<0.05; **, P<0.01; ****, P<0.0001.

To identify the YAP^High^ CFs sending signals to the other cells types, we performed ligand receptor analysis (CellChat)^19^ using the snRNA-seq dataset. Several ligand-receptor pairs were identified (Fig. 3C and Extended data Fig. 5) and further validated, including colony stimulating factor 1 (CSF1), bone morphogenesis protein 4 (BMP4), Tumor Growth Factor beta 2 (TGFB2), and Jagged 1 (JAG1) (Extended Data Fig. 6). We further focused on CSF1 receptor (CSF1R) signaling because it has been shown to regulates macrophage survival and proliferation^20^ and CSF1 was recently reported to be a direct YAP target^21^. To determine whether YAP activation in *Lats1/2*CKO CFs upregulates Csf1 expression and secretion to promote macrophage survival and proliferation, we examined *Csf1* and *Csf1r* expression in the ST dataset. We found that *Csf1* and *Csf1r* were both highly upregulated in the RA of *Lats1/*2*^ΔCF^* mice compared with control mice (Fig. 3D) and were highly colocalized (Fig. 3E). Next, we validated *Csf1* expression in the RA by using in situ hybridization and found that *Csf1* expression was upregulated in Hippo-deficient CFs compared to the control CFs (Fig. 3F, G). In addition, when macrophages were labelled with CSF1R antibody, we found that macrophage were sparse in the control RA (Fig. 3H, left panel) but substantially more abundant in the *Lats1/*2*^ΔCF^* RA (Fig. 3H, right panel), similar to what we found in the snRNAseq (Fig. 1I) and ST datasets (Fig. 1L). To determine whether these macrophages were proliferating, we performed pulse-chase EdU labeling and found that the percentage of EdU-positive macrophages was significantly increased in the *Lats1/*2*^ΔCF^* RA compared with the control RA (Fig. 3I). Because the *Lats1/*2*^ΔCF^* carry a Cre reporter, *Rosa26^Sun1-GFP^*, we labeled *Tcf21^iCre^*lineage cells by using a GFP antibody (Fig. 3H, white). Indeed, most of the RA *Lats1/*2*^ΔCF^* CFs were surrounded by macrophages.

Finally, we tested the functional role of Csf1 signaling in the macrophage expansion induced by Hippo-deficient CFs by treating the *Lats1/*2*^ΔCF^* mice with the CSF1R inhibitor, GW2580 (Fig. 3J). Compared with vehicle-treated *Lats1/*2*^ΔCF^* mice, inhibitor-treated mice showed a significantly reduced number of macrophages (Fig. 3K,L), indicating that CSF1 signaling is required for macrophage expansion in this context. Together, these results suggest that Hippo-deficient CFs secrete CSF1 to activate CSF1 signaling in macrophages, in term promoting macrophage expansion.

### Macrophages promote Hippo-deficient cardiac fibroblast proliferation and osteochondroprogenitor differentiation during fibrosis

Macrophages interact with fibroblasts during tissue repair and fibrosis in different organs^22^. Nonetheless, no injury was given in our *Lats1/*2*^ΔCF^* mice and little is known about the function of macrophages during homeostasis in vivo. To characterize the role macrophages in fibrosis induced by Hippo-deficient CFs, we inhibited macrophage expansion using the Csf1r inhibitor and examined fibrosis by staining with Sirius red. Strikingly, the fibrotic area was significantly reduced in the RA of *Lats1/*2*^ΔCF^* mice treated with Csf1r inhibitor compared with vehicle-treated *Lats1/*2*^ΔCF^* RA (Fig. 4A,B), suggesting that macrophage expansion promoted fibrosis in this context.

**Fig. 4.**
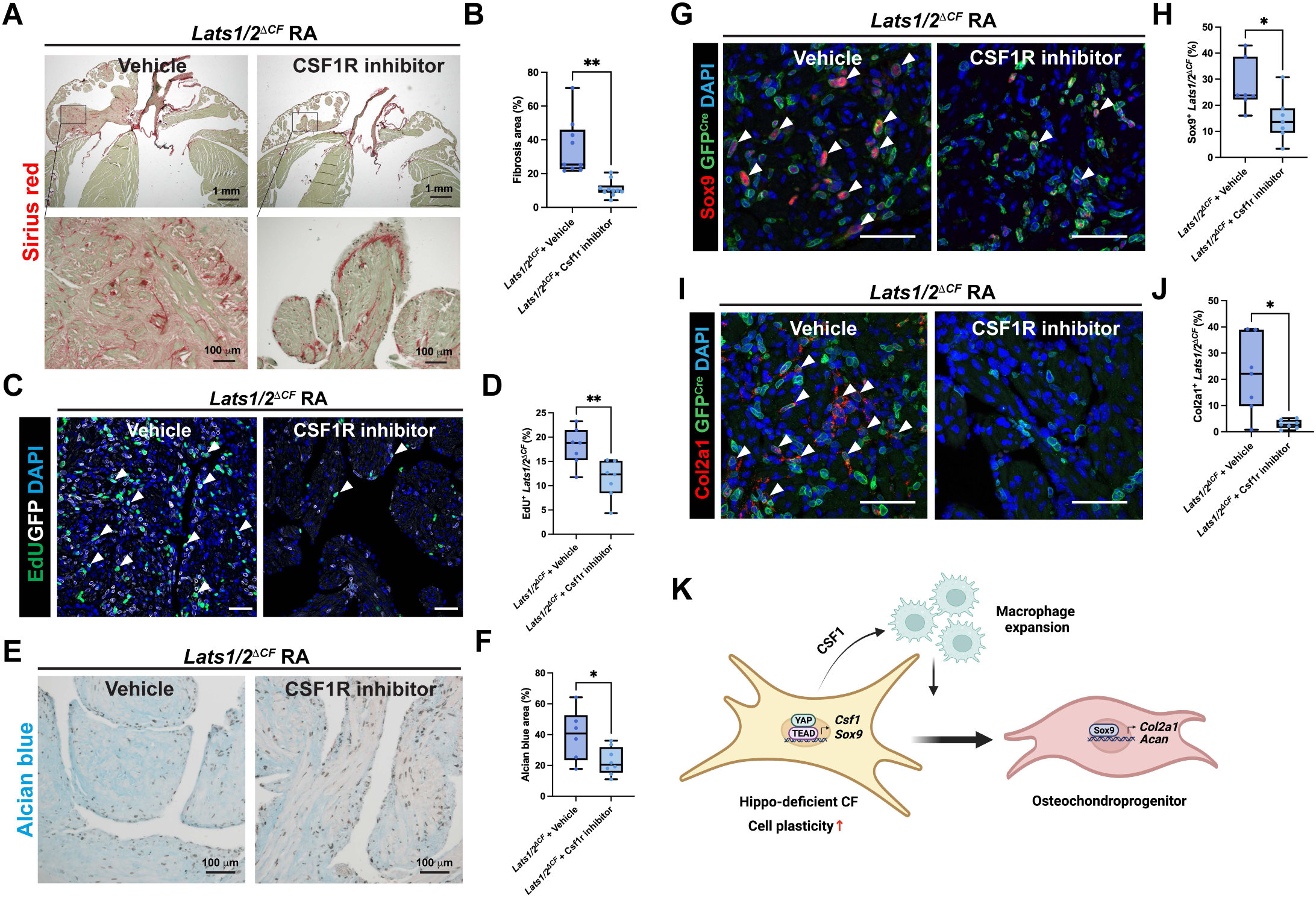
Macrophages promote Hippo-deficient cardiac fibroblasts to differentiate into osteochondroprogenitors. (A) Sirius red-stained histologic sections from the RA of *Lats1/2^ΔCF^*mice treated with vehicle or CSF1R inhibitor. Lower panels: zoomed-in images of the boxed regions in the upper panels. (B) Quantification of collagen area over tissue area. Vehicle, n=9. CSF1R inhibitor, n=10. Welch’s t-test was used for comparison. (C) Proliferating CFs in the RA of *Lats1/2^ΔCF^* mice treated with vehicle or CSF1R inhibitor were labeled with EdU staining (green). Total *Lats1/2* conditional knockout (CKO) CFs were marked by GFP (white). Scale bar, 50 μm. (D) Quantification of the EdU-positive *Lats1/2*CKO CFs in the RA of mice treated with vehicle (n=7) or CSF1R inhibitor (n=7). Welch’s test was used for comparison. (E) Alcian blue/nuclear fast red staining of sections from the RA of *Lats1/2^ΔCF^* mice treated with CSF1R inhibitor or vehicle. (F) Quantification of the Alcian blue-stained area (blue) over the total tissue area in (E). Vehicle, n=6. CSF1R inhibitor, n=8. An unpaired t-test was used for comparison. (G) Immunostaining of SOX9 (red) and GFP (green, *Lats1/2*CKO CFs) in sections from the RA of *Lats1/2^ΔCF^* mice treated with vehicle or CSF1R inhibitor. White arrowheads point to examples of SOX9-positive *Lats1/2*CKO CFs. DAPI (blue) was used to stain nucleus. Scale bar, 50 μm. (H) Quantification of SOX9-positive cells within the *Lats1/2*CKO CF (GFP+) population in the vehicle-treated (n=7) or CSF1R inhibitor-treated group (n=7). Welch’s t-test was used for comparison. (I) Co-detections of *Col2a1* transcripts (red dots) by in situ hybridization and *Tcf21^iCre^* lineage cells by GFP immunostaining (green) in sections of RA from *Lats1/2^ΔCF^* mice treated with vehicle or CSF1R inhibitor. DAPI (blue) was used to stain nuclei. White arrowheads point to examples of *Col2a1*-positive *Lats1/2*CKO CFs. Scale bar, 50 μm. (J) Quantification of *Col2a1*-positive cells within *Lats1/2*CKO CF population in the vehicle-treated (n=7) or CSF1R inhibitor-treated group (n=7). Welch’s t-test was used for comparison. (B,D,F,H,J) *, P < 0.05; **, P < 0.01. (K) Working mode: *Lats1/2*CKO CF secreted CSF1 to promote macrophage expansion. Macrophages, in turn, promote *Lats1/2*CKO CFs to differentiate into OCPs.

Macrophages have been shown to promote fibroblast proliferation in vitro^23^. Because YAP activation in Hippo-deficient CFs drive cell proliferation in a cell-autonomous manner, we examined whether the surrounding macrophage could still affect Hippo-deficient CF proliferation. To test this, we labeled proliferating cells with EdU in the hearts of *Lats1/*2*^ΔCF^*mice treated with CSF1R inhibitor or vehicle. We quantified the percentage of EdU-positive Hippo-deficient CFs and found that the percentage of EdU-positive Hippo-deficient CFs was significantly reduced in the inhibitor-treated group compared with the vehicle-treated group (Fig. 4C,D), indicating that surrounding macrophages promote Hippo-deficient CF proliferation.

Macrophages promote chondrogenesis in vitro^24^. Therefore, we tested whether macrophages are required for OCP differentiation. We first examined whether proteoglycans are reduced in the RA of *Lats1/*2*^ΔCF^* mice treated with CSF1R inhibitor by staining with Alcian blue. After treatment with Csf1r inhibitor, the *Lats1/*2*^ΔCF^* RA of the CSF1R inhibitor-treated group showed a significant reductions in Alcian blue staining compared with the *Lats1/*2*^ΔCF^* RA treated with vehicle (Fig. 4E,F), suggesting that macrophages regulate OCP differentiation. To further corroborate this finding, we examined SOX9 protein level and *Col2a1* mRNA expression in the RA of *Lats1/*2*^ΔCF^* mice treated with CSF1R inhibitor or vehicle. The expression of both SOX9 (Fig. 4G,H) and *Col2a1* (Fig. 4I,J) expression was significantly reduced in *Lats1/*2*^ΔCF^* RA treated with Csf1r inhibitor compared with vehicle-treated. These results suggest that macrophages are required for CFs to differentiate into OCPs in *Lats1/*2*^ΔCF^* RA (Fig. 4K).

## Discussion

In this study, we discovered that deleting *Lats1/2* in the CFs of mice induces OCP differentiation. Three lines of evidence support this finding. First, from the ST dataset, chondrocyte and osteoblast markers were enriched in the *Lats1/*2*^ΔCF^* RA. Second, from the snRNAseq dataset, the Hippo-deficient CF subcluster showed high expression of chondrocyte and osteoblast markers. Finally, the expressions of proteoglycans, SOX9 protein and *Col2a1* transcript was all significantly increased in the RA of *Lats1/*2*^ΔCF^* mice compared with control mice. Our study further showed that macrophage expansion induced by Hippo-deficient CFs further promote fibroblast proliferation, fibrosis, and OCP differentiation.

In *Lats1/*2*^ΔCF^* mice, we observed a much stronger inflammation and fibrosis phenotype in the RA than in the LA. However, when we compared *Tcf21^iCre^* efficiency based on Cre reporters, we found that Cre activity was similar between the LA and RA. Moreover, when we compared the knockout efficiency of *Lats1/2* between the LA and RA by using real-time qPCR, no significant difference was observed. Thus, we excluded the possibility that the weaker phenotype in the *Lats1/*2*^ΔCF^* LA versus the RA was due to lower Cre activity in the LA. We speculate that CFs may be different between the LA and RA, which has been reported in human hearts on the basis of single-nucleus transcriptomics data^25,26^. Because the Hippo pathway in CFs plays a key role in organ fibrosis^27,28,29^, understanding the key differences between LA and RA CFs that affect Hippo pathway should provide insight into fibroblast biology and treatment designs for fibrotic diseases.

CFs increase cell plasticity after injuries or under stress^2,30,31^. Lineage-tracing experiments in mice showed that, upon injury, resting CFs differentiated into activated fibroblasts, myofibroblasts^32^ and matrifibrocytes^33^, in addition to acquired osteogenic fate^6^ and adipogenic fate^5^. Nonetheless, the differentiation potential of CFs during homeostasis remains unclear. Here, we showed that in the absence of cardiac insult, YAP activation in CFs drives OCP differentiation. This raises the question of how YAP promote cell plasticity and OCP differentiation. Because SOX9 is a key transcriptional factor during chondrogenesis^34,35^, it is possible that YAP directly upregulates *Sox9,* similar to that reported in a cancer context^36^. Interestingly, YAP also inhibits SOX9-induced chondrogenic program in neural crest cells^37^. Thus, it is possible that YAP increases SOX9 protein but simultaneously inhibits its activation, which pois CF into this OCP cell state. In the current study, we also showed that macrophages are required for the expression of SOX9 protein in RA *Lats1/2*CKO CFs, suggesting the non-cell autonomous regulation of SOX9 by macrophage, presumably through macrophage-secreting factors. Future studies are warranted to understand the detail underlying YAP-SOX9 regulation and how macrophages affect CF cell state changes.

GWAS have shown that variants in the genes encoding Hippo pathway components are associated with human congenital heart defects and heart disease, such as septal and valve defects and arrythmia. Recently, deleting *Lats1/2* in pacemaker cells induce arrythmia in mice^38^. In current study, we also found that deleting *Lats1/2* in CFs caused arrythmia, abnormal P-wave and heart rate. Furthermore, inflammation and fibrosis are associated arrythmia^39^. For future studies, our mouse model provides an ideal system for investigating how inflammation and fibrosis affect the cardiac conduction system.

## Acknowledgments

This work is supported by National Institutes of Health grant HL 127717 (J.F.M.), National Institutes of Health grant HL 130804 (JFM), National Institutes of Health grant HL 118761 (J.F.M.), Vivian L. Smith Foundation (J.F.M.), Don McGill Gene Editing Laboratory of The Texas Heart Institute (X.L.).

Author contributions: C-R.T. and J.F.M conceptualized the project; C-R.T., J.K., X.L., P.C. were responsible for Methodology; C-R.T., J.K., X.L., P.C., R.L., F.M., M.Z., X.Z., F.G., J.S. conducted the investigation; C-R.T. wrote the original draft; C-R.T. and J.F.M reviewed and edited the manuscript; J.F.M., X.L. acquired funding. J.F.M. supervised the project.

