## Extended data for "Hippo-deficient cardiac fibroblasts differentiate into osteochondroprogenitors"

### Extended Data Fig. 1

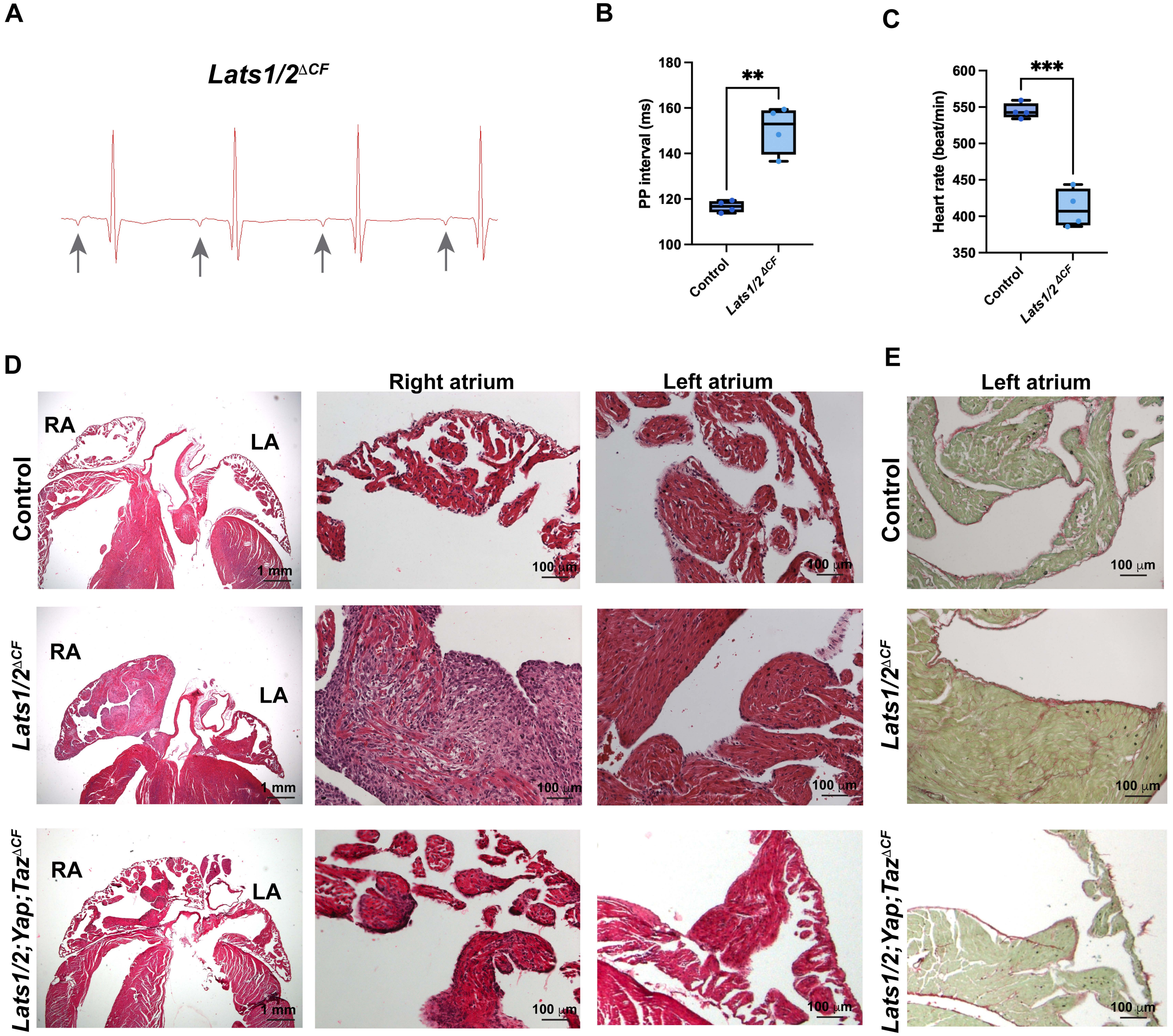

### Extended Data Fig. 2

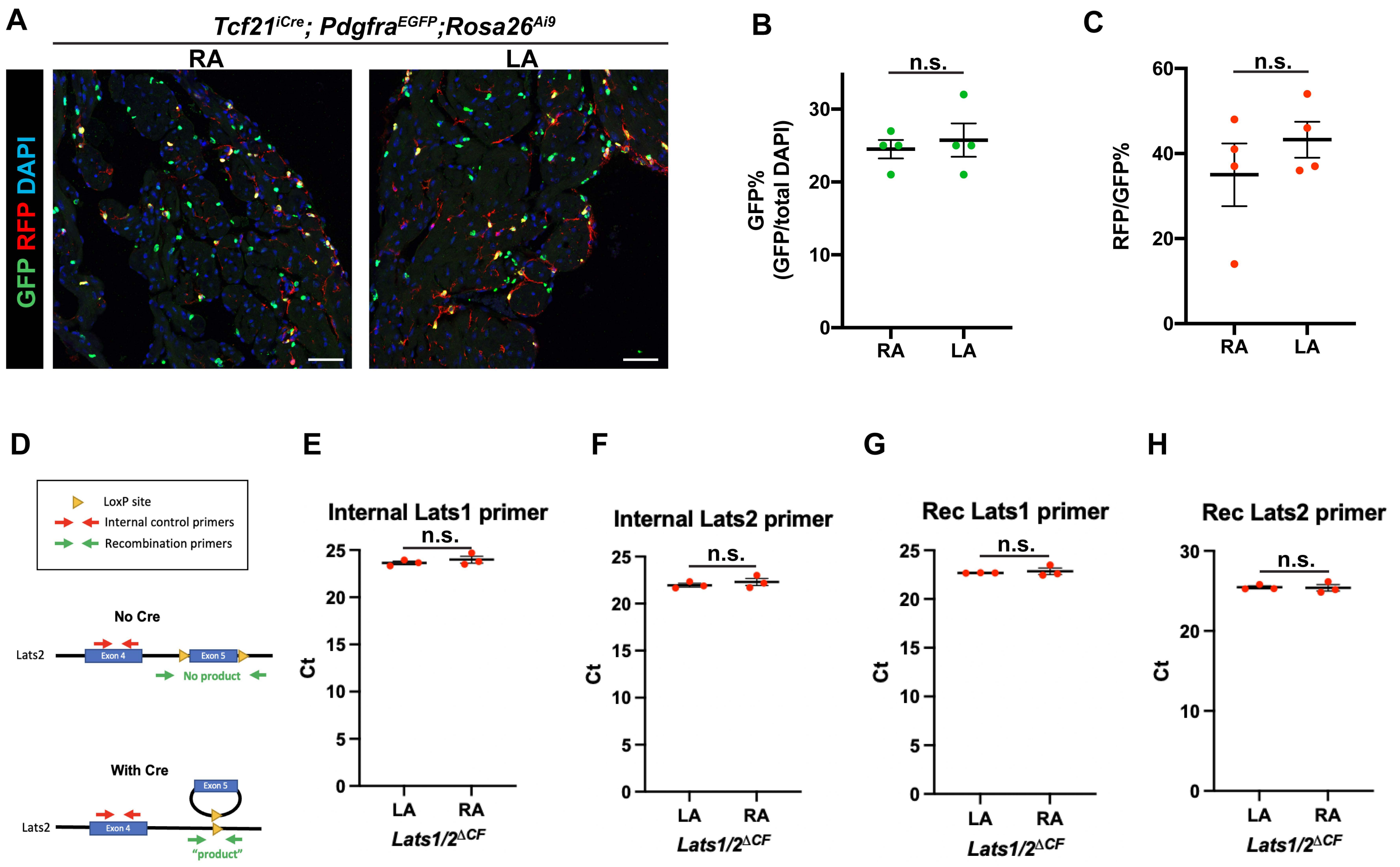

### Extended Data Fig. 3

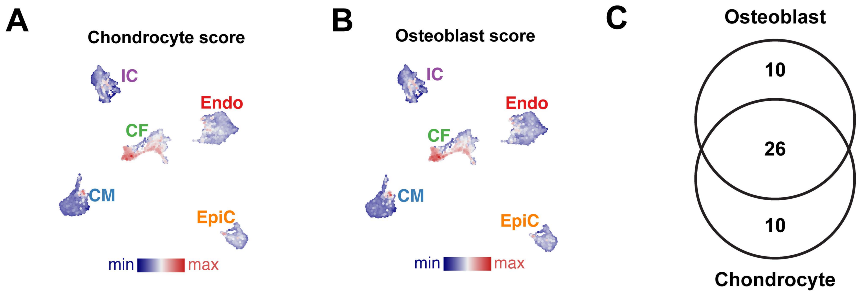

### Extended Data Fig. 4

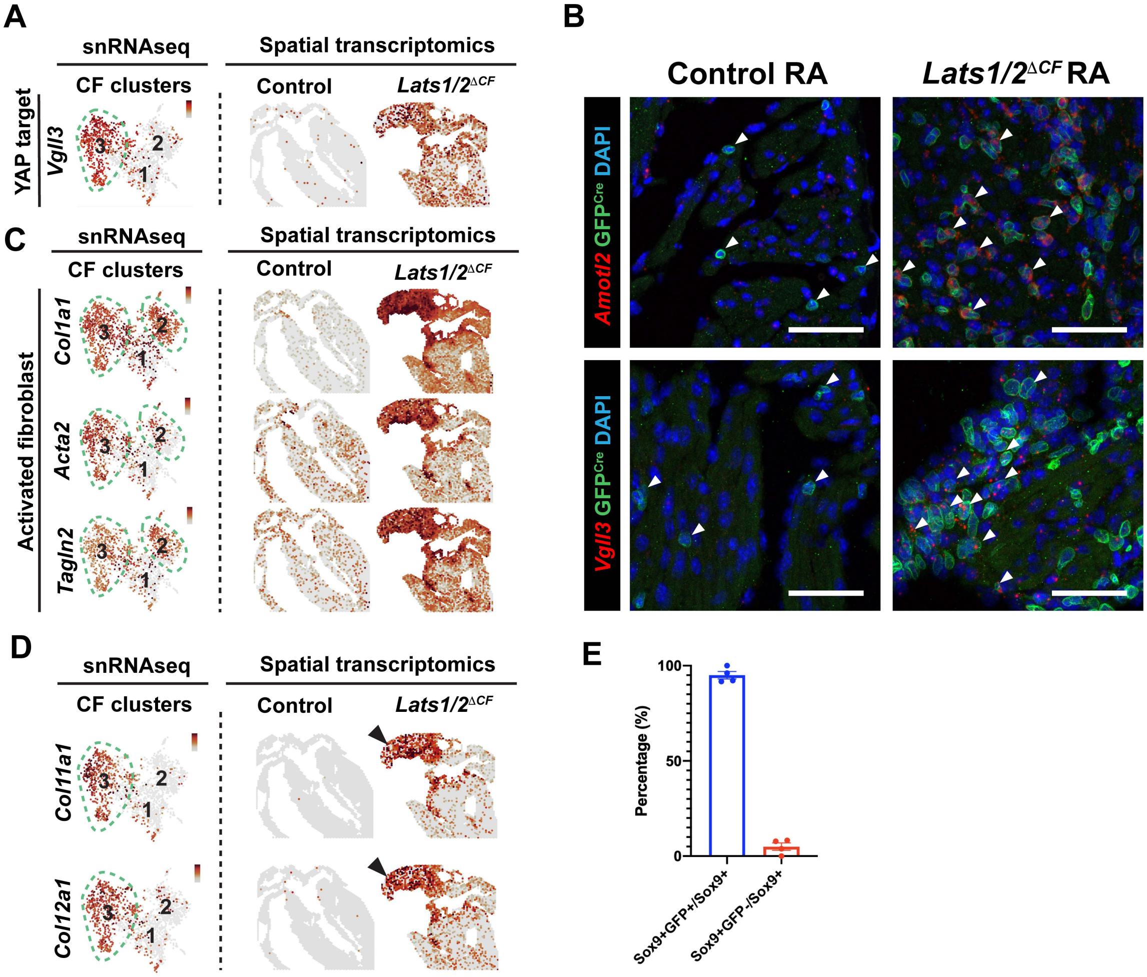

### Extended Data Fig. 5

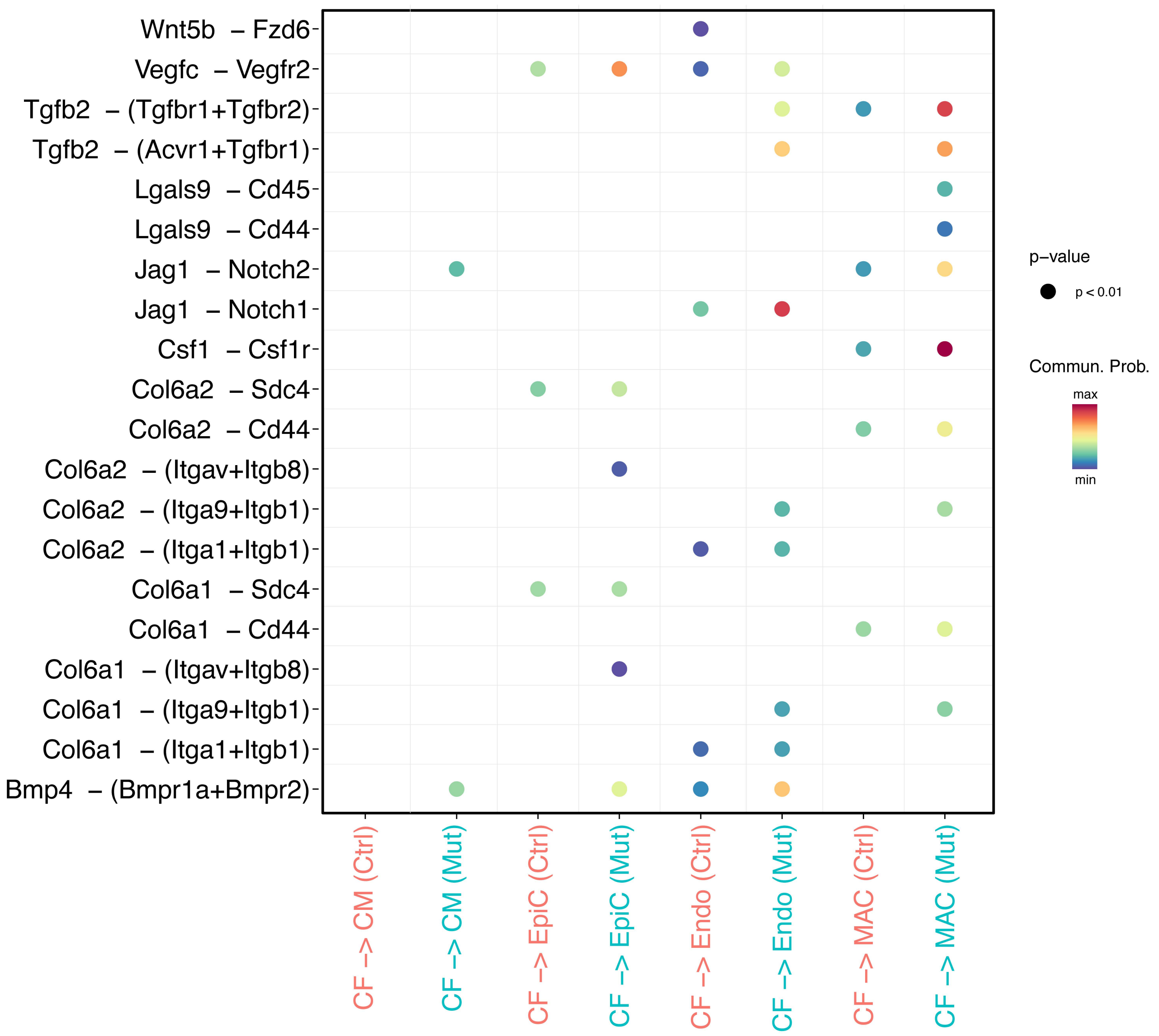

Extended Data Fig. 6

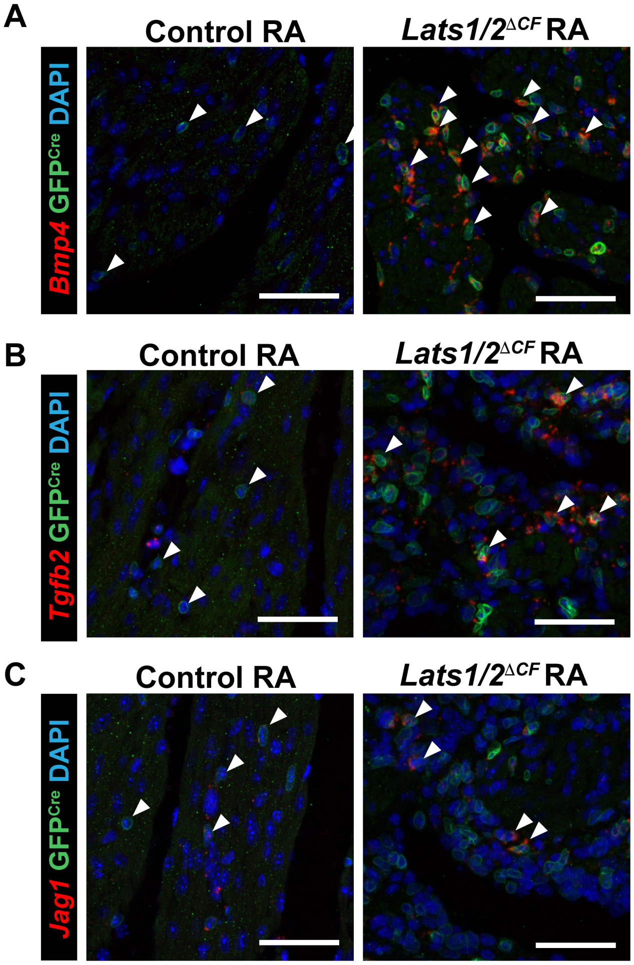
